# Novel ankyrin-repeat mutant and modifiers of a kafirin mutant improve sorghum protein digestibility

**DOI:** 10.1101/863951

**Authors:** Elisabeth Diatta-Holgate, Mitchell Tuinstra, Charles Addo-Quaye, Ndiaga Cisse, Agyemang Danquah, Pangirayi Tongoona, Eric Danquah, Clifford F. Weil

## Abstract

Sorghum is a staple food for over 500 million people in Sub-Saharan Africa and Asia, however, sorghum proteins are poorly digested when wet-cooked. Three sorghum mutants were identified in a mutagenized population of the inbred line BTx623 that showed a 23-37% increase in wet-cooked protein digestibility compared to their unmutagenized parent. Furthermore, in comparison to the known high lysine, highly digestible sorghum mutant, P721Q, these mutants had 9% more protein overall that was 10% more digestible, had 12% more lysine, as well as better seed hardness. Using bulked segregant analysis based on whole genome sequencing data, we identified unique genomic regions on chromosome 5 of each EMS mutant that are associated with the increase in protein digestibility. Analyzing shared mutations in candidate genes, the high protein digestibility phenotype in one mutant is linked to a point mutation in a novel, ankyrin repeat protein. In another, the increase is associated with a mutation in a kafirin gene and suggests novel genetic modifiers. This study provides material and molecular markers that can be used to enhance sorghum nutritional value, contribute to fighting malnutrition and elucidate new roles for ankyrin-repeat proteins in plants.

**One sentence summary:** Mutations in a novel, ankyrin domain protein and genetic modifiers of a known mutation in a seed storage protein lead to increased digestibility of seed proteins in sorghum after wet cooking.

## INTRODUCTION

In many parts of Sub-Saharan Africa and South Asia, sorghum is the main cereal crop grown and consumed as a staple food. It is a relatively low-cost grain and is adapted to warm and low rainfall environments. Nonetheless, due to low digestibility of its proteins after wet cooking (Axtell *et al.*, 1981; reviewed by Duressa *et al*, 2018), sorghum needs to be supplemented in diets with other crops that are drought susceptible and/or use nitrogen less efficiently and that are often too expensive for those who rely on sorghum as a staple. Additionally, sorghum protein exhibits low quantities of the essential amino acids tryptophan and lysine, which can lead to malnutrition, especially in places where sorghum is one of the primary sources of protein (Maclean Jr *et al.*, 1981). It is therefore important to solve this problem to contribute to food security.

Previous research on sorghum led to the identification of a high lysine-highly digestible mutant named P721Q (opaque endosperm), generated from P721N (normal endosperm) (Mohan, 1975; Weaver *et al.*, 1998). Induced using the mutagen diethyl sulphate, P721Q had a G:A mutation in one of the 20 genes encoding the major seed storage protein, alpha-kafirin, located on chromosome 5; the mutation prevents the cleavage of the signal peptide from the kafirin (Wu *et al.*, 2013). However, exactly which copy of the alpha-kafirin gene has been mutated remains unclear. Kafirins are alcohol soluble prolamins, the main storage proteins in sorghum endosperm, and are found inside round protein bodies (Shull *et al.*, 1991). P721Q had unusual, highly invaginated protein bodies and, although P721Q had 60% more lysine (Mohan, 1975) and 25% more digestible proteins than its parent line (Weaver *et al.*, 1998), it had soft seeds with floury endosperm, making it susceptible to bird attacks and damage during handling, and was not very suitable for some African meals, thus limiting consumer acceptance. To achieve higher digestible protein with better seed quality, an ethyl methane sulfonate (EMS) mutagenized population of BTx623 was screened using a modified protein digestibility (PD) assay (see Methods) (Mertz *et al.*, 1984; Aboubacar *et al.*, 2003). Three highly digestible mutants, SbEMS1613, SbEMS1227, and SbEMS3324 were identified (Massafaro, 2015).

Here we characterize these EMS mutants genetically, assess grain quality, investigate the mechanisms underlying the increase in protein digestibility and develop molecular markers that could facilitate rapid introgression of the improved digestibility trait into locally-adapted sorghum lines. Using bulked segregant analysis, we have identified candidate genes that could be responsible for the increased digestibility in the EMS mutants. We then created half-sib progenies that only had in common the highly digestible mutant as parent and compared their genomes for shared mutations. This approach is a fast, efficient and cost-effective method of identifying causative genes in a mutant background.

## RESULTS

### Protein digestibility of the mutants

We assessed the difference in protein digestibility between sorghum mutants and their unmutagenized parents under both uncooked and wet-cooked conditions. Protein digestibility (PD) was determined using a modified PD assay where samples were wet-milled, then either cooked or not and digested with pepsin to mimic typical sorghum usage as food or feed. The EMS mutants were between 25 and 37% more digestible than the unmutagenized control after wet cooking (Figure 1). In particular, SbEMS3324 showed a 15% improvement in protein digestibility over P721Q.

**Figure 1.**
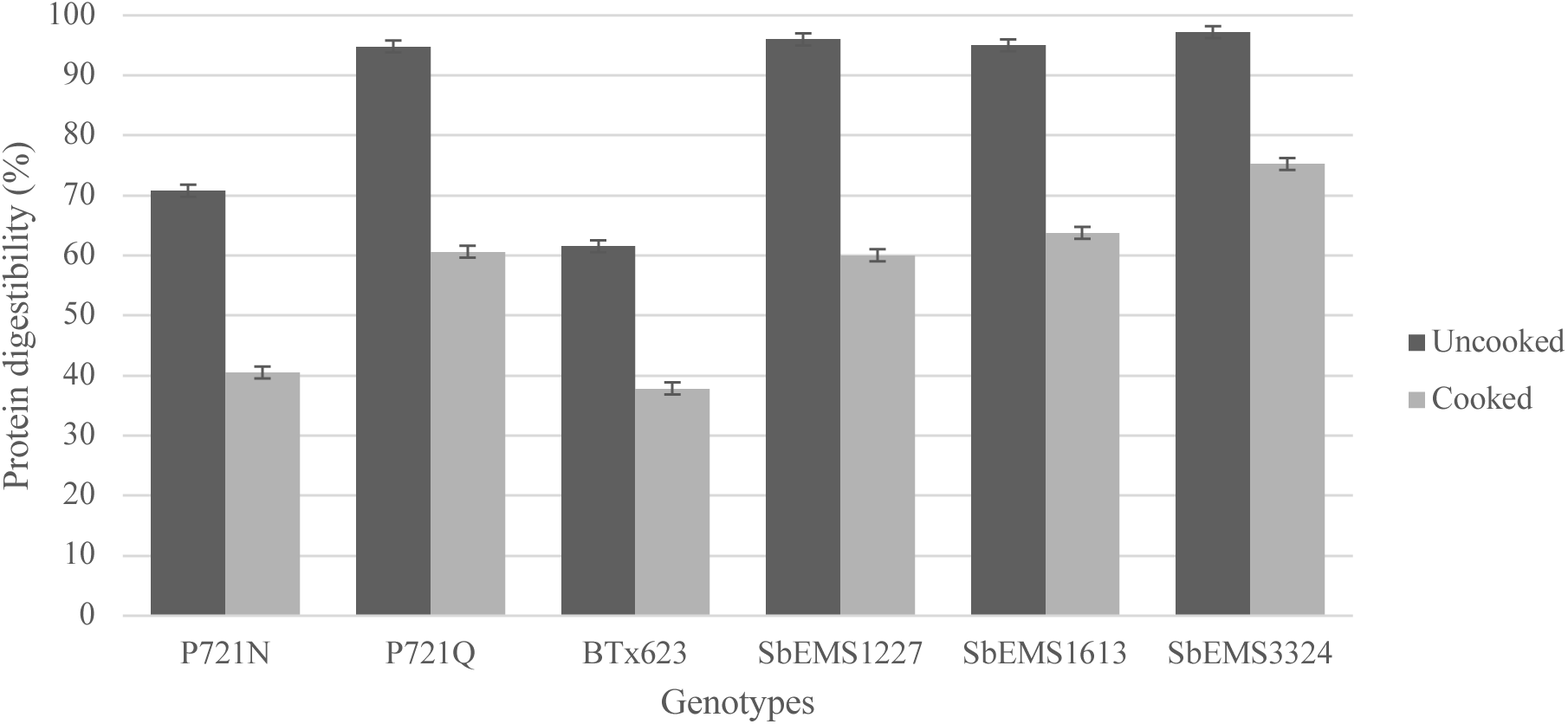
Comparison of average protein digestibility. Error bars show standard error.

### Protein body morphology of the EMS mutants

Given the altered morphology of protein bodies reported for P721Q (Oria et al, 2000), and the hypothesis that this change in structure may relate to increased digestibility, we examined the protein bodies of the three EMS mutants (SbEMS1227, SbEMS1613, SbEMS3324) in comparison to their progenitor (BTx623). All 4 lines were grown in the greenhouse and seeds, collected at 30 days after half bloom, were fixed, sectioned and examined by transmission electron microscopy (Figure 2). The protein bodies of the wild type BTx623 were round, as expected, and interestingly, the protein bodies of the highly digestible SbEMS1613 mutant were also round (Figures 2A and 2B). In contrast, SbEMS1227 had highly invaginated protein bodies similar to those reported for P721Q (Figure 2C) and SbEMS3324 had a mixture of protein body morphologies, some round and some with an intermediate level of invagination (Figure 2D). Taken together, these data suggest that protein body morphology is not the sole explanation for increased digestibility.

**Figure 2.**
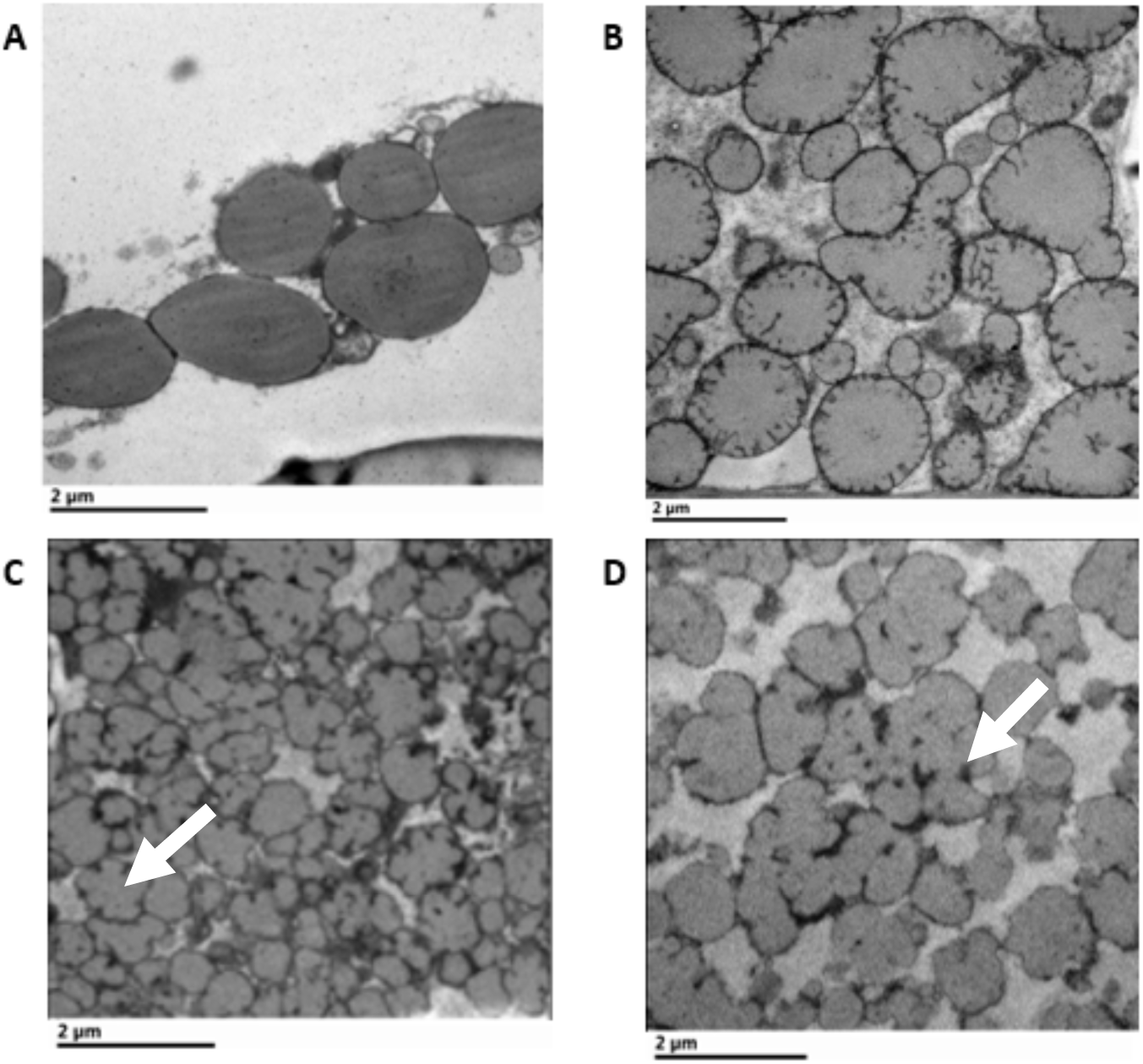
Transmission electron microscopy images of sorghum seed protein bodies. Invaginated protein bodies indicated by arrows. BTx623 (A) and highly digestible mutants SbEMS1613 (B), SbEMS1227 (C), and SbEMS3324 (D).

### Bulk Segregant Analysis

The EMS mutations were mapped by bulked segregant analysis (BSA). To obtain the bulk samples needed, we created mapping populations by crossing each highly digestible line to a low digestibility line carrying a *brown midrib* mutation, allowing us to distinguish the F1 hybrid. These F1 were then selfed, F2 plants grown, sampled for DNA and selfed again, and triplicate pairs of F3 seed assessed for digestibility. Plants for which all three seed samples were highly digestible compared to controls (consistently ≥ 60% protein digestibility) were scored as homozygous mutant, plants where all three samples were comparable to the unmutagenized progenitor were scored as homozygous wild-type and plants with mixed results were scored as heterozygous and not included in the bulked segregant analysis. Two additional samples from all the homozygous high and homozygous low candidates were tested to confirm digestibility status.

Protein digestibility in the mapping populations for SbEMS1613 (Figure 3a), SbEMS3324 (Figure 3b) and SbEMS1227 (data not shown) followed a bimodal distribution showing two distinct groups, each with means similar to the low and high digestibility parents for each cross. Each F2 fit a 3:1 segregation ratio (p < 0.05), consistent with a trait controlled by a single, recessive mutation in each case. Interestingly, the highly digestible bulk for the cross of SbEMS3324 had a mean digestibility even higher than the SbEMS3324 parent, a transgressive phenotype suggesting the possible presence of a genetic modifier.

**Figure 3.**
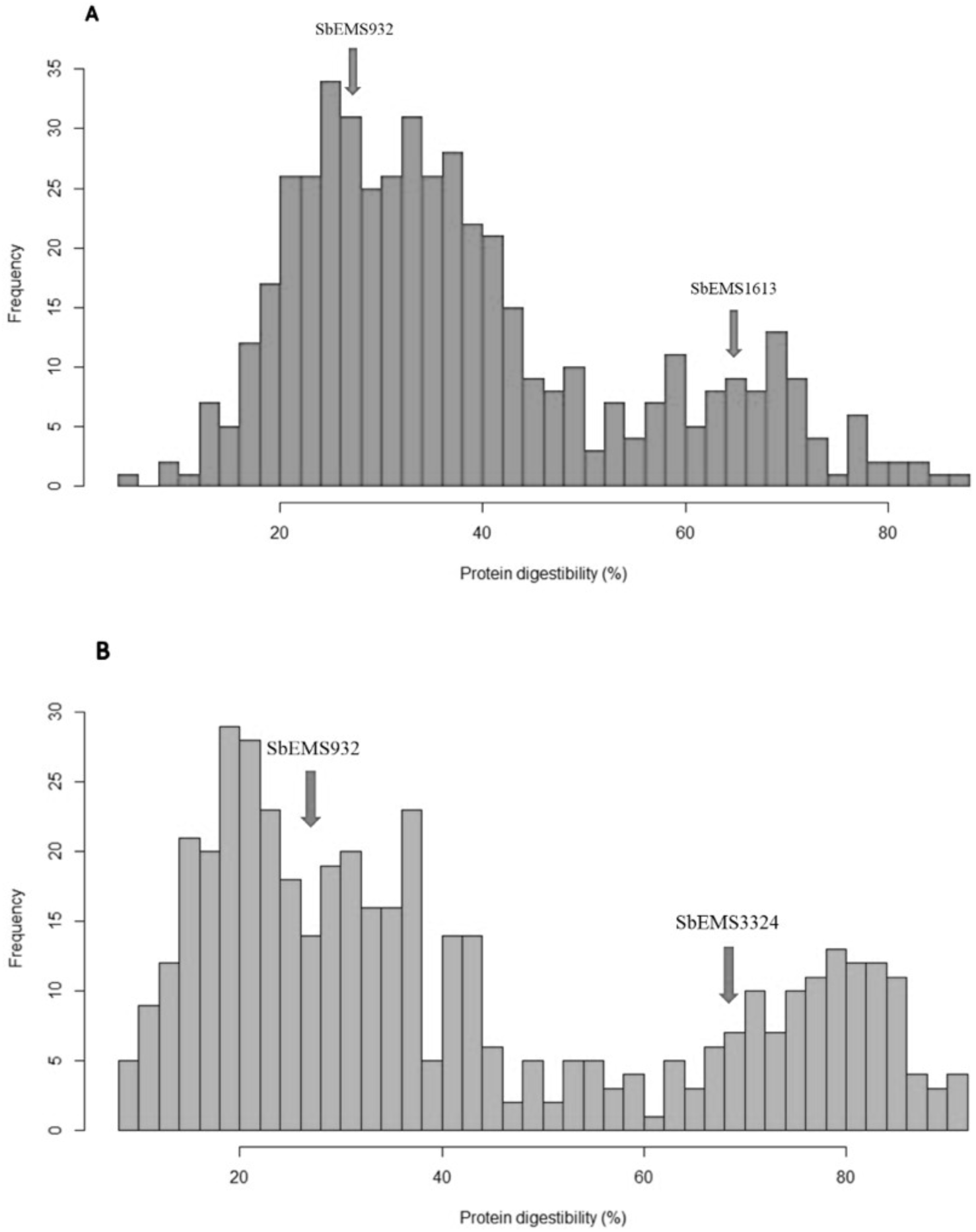
Frequency distribution of the corrected protein digestibility values (BLUEs) for mapping populations of SbEMS1613 (A) and SbEMS3324 (B). Digestibility of the low (SbEMS932) and high (SbEMS1613 and SbEMS3324) parents indicated by arrows.

We were particularly interested in mutants likely to differ from the P721Q mutant reported previously, so focused on SbEMS1613 and SbEMS3324 because of the differences in their protein body morphologies as compared to P721Q. To identify the causative allele(s) for each EMS mutant, we performed whole genome sequencing on bulked DNA from homozygous high and homozygous low plants for each mutant. At most loci, we expect a 1:1 ratio of each parental allele in the bulk progenies. However, at loci linked to the mutation responsible for the difference in protein digestibility between the high and the low bulk, we expect the allele frequency in the high bulk to be shifted in favor of the allele(s) from the highly digestible parent and, ideally, to cosegregate.

### SbEMS1613

High and low digestibility groups used for BSA were made of 50 entries each for SbEMS1613; the high group ranged from 60.12 to 86.8% digestibility and the low group from 11.96 to 30.08%. Equal amounts of leaf tissue for each plant in a bulk were combined, DNA was extracted and whole genome sequenced and a total of 104,253 and 102,663 SNPs were obtained in the high digestibility and low digestibility bulks, respectively. The difference in SNP allele frequencies compared to the BTx623 genome for the high and the low bulks are plotted in Figure 4A.

**Figure 4.**
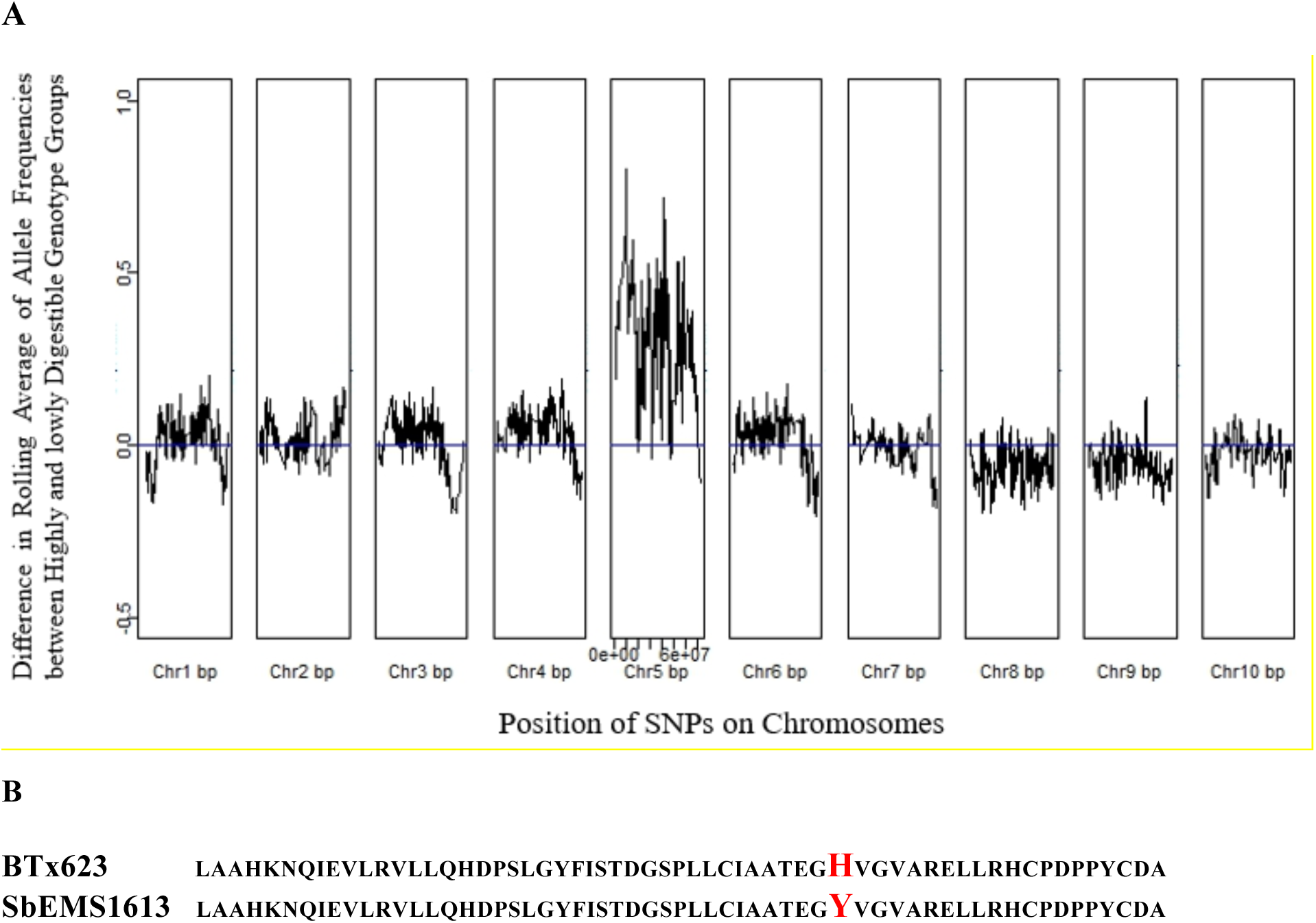
Bulked segregant analysis in F_2_ segregants of SbEMS932 × SbEMS1613 mapping population. A) Comparison between allele frequencies of high and low bulks B) Partial comparison of protein sequences encoded by Sobic.005G083340 showing mutation in one of the ankyrin repeats in red.

A strong association with high protein digestibility was detected on chromosome 5. Indeed, all of chromosome 5 seems to be biased towards the high digestibility allele(s), with a strong peak between 39 Mb and 43 Mb and a minor peak at 11-12 Mb. There were no SNPs found within annotated genes in the region under the major peak. However, a C:T mutation was found under the minor peak, at position 1100 within the coding region of Sobic.005G083340, an ankyrin repeat-containing protein annotated as similar to the 26S proteasome non-ATPase regulatory subunit PSMD10 (Phytozome 12). The allele frequency was equal to 1.00 in the high bulk and 0.16 in the low bulk, consistent with expectations for a recessive allele. This mutation is predicted to cause a non-synonymous missense mutation, H234Y (Fig 4B), which lies within an ankyrin domain that is conserved in orthologs of this protein found throughout the grasses.

### SbEMS3324

BSA for SbEMS3324 was performed with two bulks of 30 samples each with protein digestibility ranging from 62.7 to 91.87% for the high group and 11.39 to 30.84% for the low bulk. A total of 181,744 G:C to A:T SNPs was identified for the high digestibility bulk and 175,899 G:C to A:T SNPs for the low digestibility bulk. The difference in allele frequencies between the high and the low bulks again showed that the entire chromosome 5 was biased towards alleles in the high digestibility bulk, but in this case, there was one major peak around 67 Mb (Figure 5A). The only gene with a G:C to A:T mutation in this region encodes a kafirin protein (Sobic.005G189000). This missense mutation is predicted to cause an A21T substitution at the cleavage site of the signal peptide of the kafirin (Figure 5B) and, interestingly, is identical to the amino acid substitution identified in P721Q. Comparison of SNPs in the surrounding DNA sequence with those in P721Q verified that this is a new allele of BTx623 and not a contaminant (data not shown).

**Figure 5.**
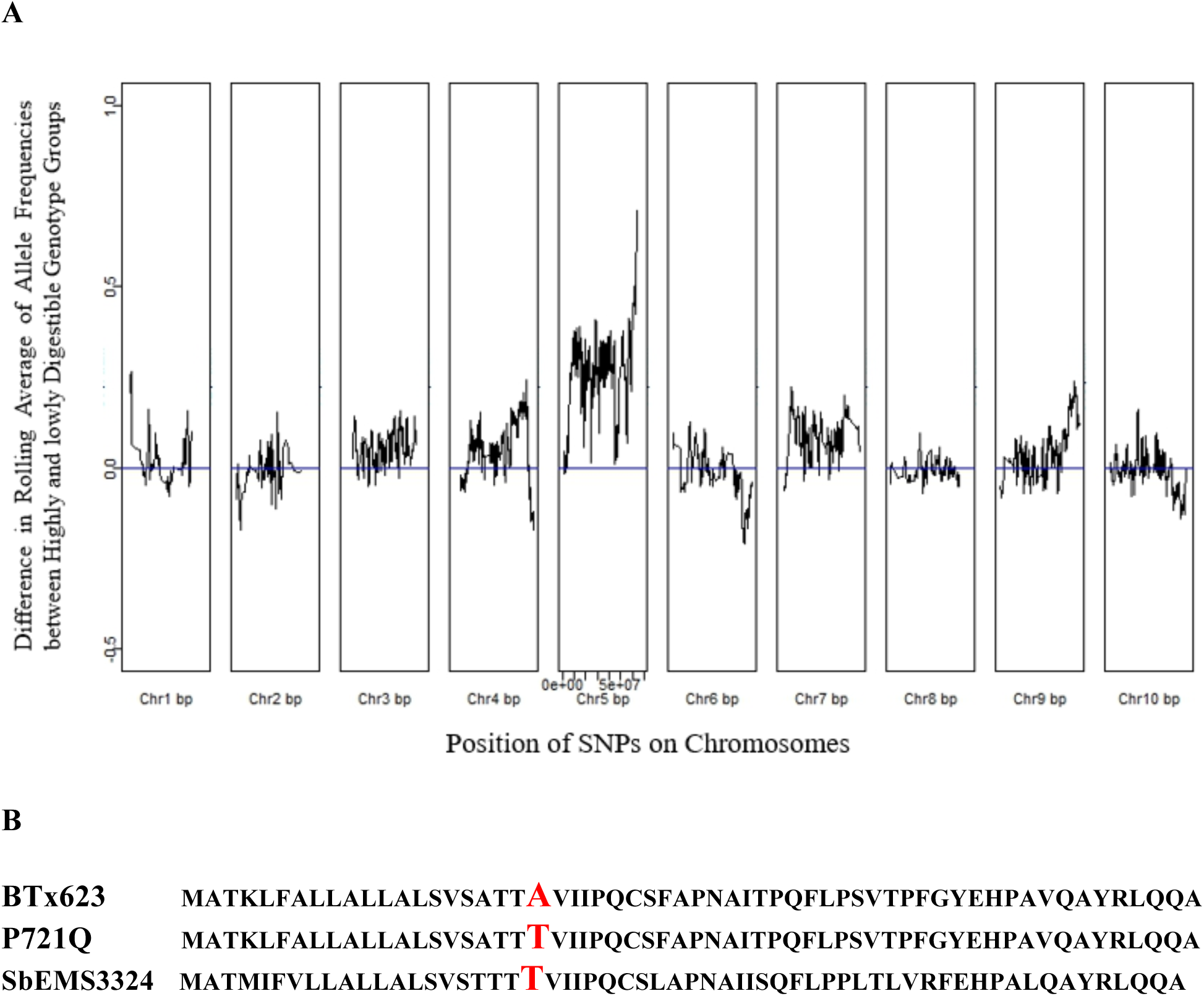
Bulked segregant analysis in F_2_ segregants of SbEMS932 × SbEMS3324 mapping population. A) Comparison between allele frequencies of the high and low bulks. B) Comparison of protein sequences for *Sobic.005G189000*

### SNP validation

In both BSA datasets, the mutations described above cosegregated perfectly with the phenotype, present at 100% in the high digestibility bulks. We crossed SBEMS1613 to three and SbEMS3324 to four other inbred lines of sorghum (see Materials and Methods), none of which carried the high digestibility alleles at these positions, selfed, sampled F2 leaf tissue and tested F3 seed from each of the crosses in triplicates. In both cases, all F2 lines identified as homozygous highly digestible (n=36 and n=35 respectively) had the mutations in either the ankyrin repeat protein or the kafirin gene (data not shown), continuing to cosegregate perfectly with the phenotype. None of the homozygous low digestibility plants had a mutation in the candidate genes.

### Protein, lysine and tryptophan contents of highly digestible sorghum mutants

The P721Q mutant was originally isolated as a high lysine variety and only later discovered to have improved digestibility (Mohan 1975). We therefore tested our mutant seeds for changes in either total protein content or increased proportions of the essential amino acids lysine and tryptophan compared to the BTx623 progenitor (Table 1). In both SbEMS1613 and SbEMS3324, crude protein and the proportion of lysine and tryptophan are higher.

**Table 1:** Crude protein, total lysine and tryptophan content of wild type and sorghum mutants.

| Sample ID | Crude protein<br>(g/100 g of samples) | Total lysine<br>(g/100 g of samples) | Tryptophan<br>(g/100 g of samples) |
| --- | --- | --- | --- |
| BTx623 | 10.66 | 0.24 | 0.09 |
| SbEMS1613 | 13.38 | 0.36 | 0.12 |
| SbEMS3324 | 14.84 | 0.48 | 0.14 |

### Seed diameter of EMS mutants

An important consideration in whether modifications to grain are adopted widely is whether they impact yield. We therefore assessed the seed size of the mutants and their progenitors, measured as diameter of mature seed (Table 2). While there is a small but statistically significant decrease in seed size comparing the EMS mutants to the BTx623 progenitor, the mutants are still larger than the P721Q seed and may therefore represent a better yielding, highly digestible alternative.

**Table 2:** Seed diameter of sorghum mutants and their wild type parents.

| Genotypes | Seed diameter (mm) |
| --- | --- |
| BTx623 | 3.98 a |
| P721N | 3.86 b |
| SbEMS3324 | 3.83 b |
| SbEMS1613 | 3.77 c |
| P721Q | 3.64 d |

### Seed hardness of EMS mutants

Another characteristic of the known highly digestible mutant P721Q is that it produces a soft, floury endosperm, leaving it susceptible to bird and insect, as well as handling damage, limiting its utility and necessitating additional breeding to find genetic modifiers that improve seed hardness. Comparing the mechanical strength of seeds, wild type sorghum lines P721N and BTx623 had a similar level of seed hardness and the two kafirin mutants (P721Q and SbEMS3324) had similarly softer seed (Figure 6). However, seed of the highly digestible SbEMS1613 mutant was significantly harder than the kafirin mutants (17%; Figure 6).

**Figure 6.**
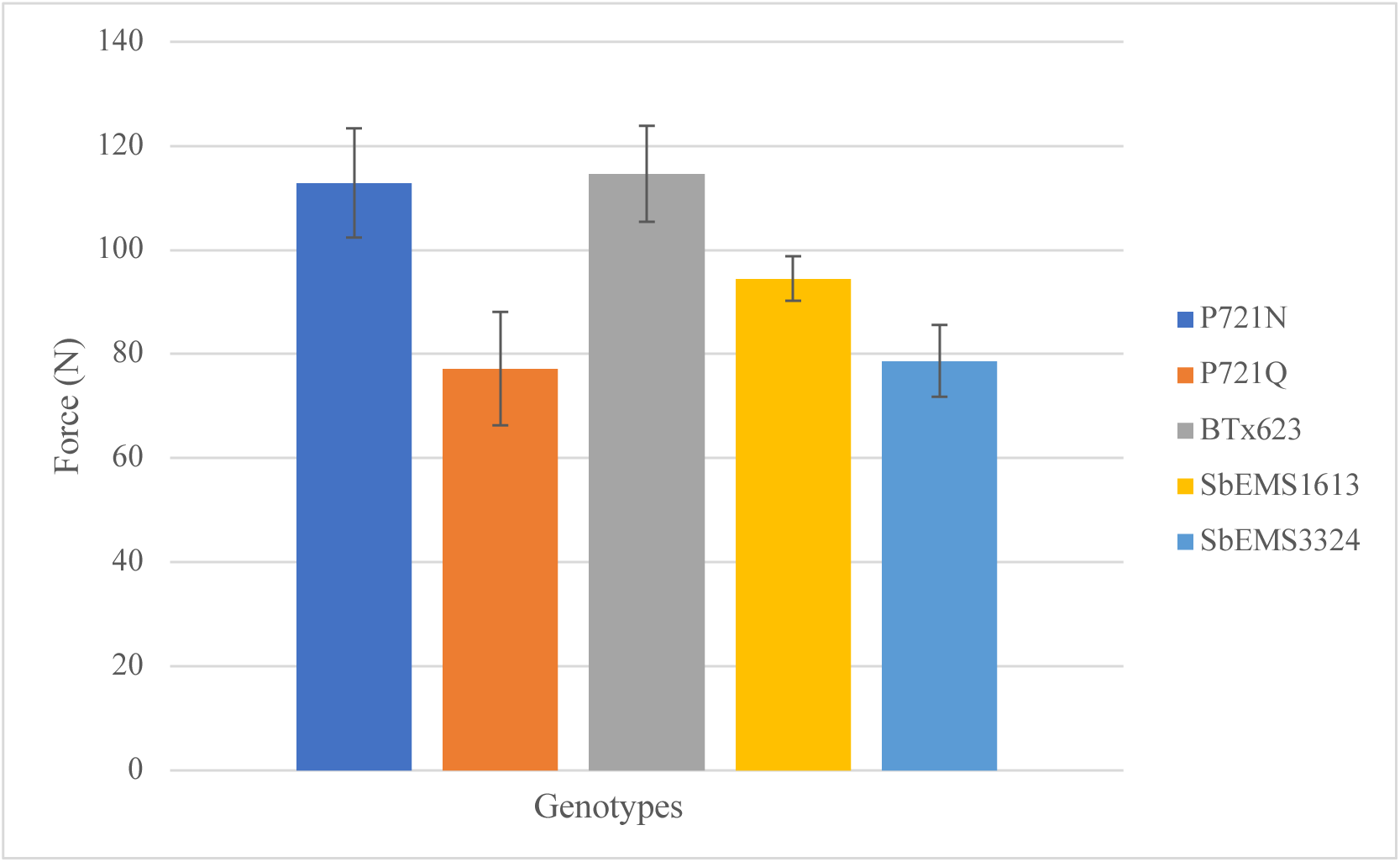
Seed hardness of mutants P721Q, SbEMS1613 and SbEMS3324 compared to their wild type parents P721N and BTx623, respectively. Error bars show standard error.

## DISCUSSION

We have identified sorghum EMS mutants with significantly higher protein digestibility than their progenitor, and these mutants also have higher crude protein content, lysine and tryptophan than their wild type parent BTx623. The increase in total lysine could be a result of a reduction in lysine-poor prolamins along with an increase in more lysine-rich albumins, globulins and glutelins (Singh and Axtell, 1973; Wu *et al.*, 2013), or could be an increase in free-lysine.

The most highly digestible mutant described here, SbEMS3324 appears to carry exactly the same mutation in a kafirin-encoding gene as P721Q, and is similar in seed hardness and endosperm texture; however, it is significantly more digestible in side-by-side tests, has significantly larger seeds and has much less invagination to its protein bodies, suggesting that there are likely to be important genetic modifiers of this allele in BTx623 as compared to P721N that have yet to be explored. Another of our mutants, SbEMS1613, while still softer than its unmutagenized progenitor, had significantly harder seed than either P721Q or SbEMS3324 and carries a mutation that does not map to a kafirin gene, indicating that there are at least two mechanisms for achieving increased digestibility. Sorghum seed hardness is very important in areas where birds attack the crop, where the seed are susceptible to grain mold and where the seed are subjected to hand or industrial processing of the whole grain before end use (for example, removing the pericarp (dehulling) before cooking). At this point, the mechanistic reasons for decrease in seed size and hardness in these mutants remain unclear.

The decrease in sorghum protein digestibility when wet-cooked compared to other cereal grains is known to be influenced by a variety of factors including protein crosslinking, endosperm structure, and the presence of antinutrients such as tannins, among others (Maclean Jr *et al.*, 1981; Duodu *et al.*, 2003). In P721Q, the increase in protein digestibility was hypothesized to be associated with its highly invaginated protein bodies with the idea that the more readily accessible the kafirin proteins are, the more digestible will be the genotype. However, in this study, among the three highly digestible EMS mutants from BTx623, only SbEMS1227 displayed highly invaginated protein bodies. SbEMS3324 had a mixture of protein body phenotypes, while SbEMS1613 showed round protein bodies similar to the wild type. Therefore, while the invaginated protein body phenotype is associated with an increase in digestibility, our data suggest there are other explanations as well.

The increase in digestible proteins in SbEMS1613 after wet cooking maps to a region containing a gene encoding an ankyrin repeat protein on chromosome 5 (Sobic.005G083340) we are naming *high digestibility 1 (hdg1)*. A recessive mutation, which we are calling *hdg1-1*, that is consistent with an EMS-induced change and that would cause an H234Y missense substitution altering one of the ankyrin repeats in this protein cosegregates with the mutant phenotype. The presence of the homozygous *hdg1-1* mutant allele in all highly digestible F3 progeny from three additional crosses to other sorghum inbreds as well as the samples from our mapping populations, strongly suggests that the increase in digestibility is tightly linked to the *hdg1-1* mutation. While a second allele of *HDG1* has not yet been linked to increased digestibility, it is interesting to consider that the kafirin genes in SbEMS1613 are all nonmutant (data not shown), indicating that there are multiple ways to increase wet-cooked sorghum digestibility genetically, some of which retain better seed hardness (Fig 6). Based solely on sequence similarity, *HDG1* has been annotated as similar to the non-ATPase regulatory subunit 10 of the 26S proteasome complex, also an ankyrin repeat protein; however, BLAST searches of the sorghum genome with all the known plant genes for regulatory subunits of the 26S complex fail to identify Sobic.005G083340 as a match and we therefore suggest *HDG1* encodes some other ankyrin repeat protein of as yet unknown function. It is hypothesized that wet cooking induces protein cross linking and disulfide bond formation in sorghum protein bodies and that this is the reason for poor protein digestibility (Duodu *et al.*, 2002). Whether a mutation in an ankyrin repeat, which is often involved in protein-protein interaction, could disrupt this cross-linking will be an interesting avenue of further study.

In SbEMS3324, the G:A mutation in the gene, Sobic.005G189000, causes an A21T substitution in the signal peptide of an α-kafirin, similar to the one described for P721Q (Wu et al 2013), althooughthis is an independently derived mutant. In maize, the alanine at position 21 of the analogous storage protein, α-zein, is the site of signal peptide cleavage, and a mutation in the *floury2* gene that causes an A21V substitution causes an increase in lysine content and invaginated protein bodies in maize endosperm similar to those described here (Lending and Larkins, 1992; Kim *et al.*, 2004). In previous studies the exact copy of the kafirin gene altered in P721Q remained unknown, as many kafirin genes are located on chromosome 5 and are highly similar to one another. Using bulked segregant analysis and whole genome sequencing, the exact copy mutated in SbEMS3324 was identified in this study and sequence alignment suggests that it is the same mutation occurring in P721Q. We hypothesize that the reason for SbEMS3324 having even higher digestibility than P721Q is the presence of one or more genetic modifiers that differ between BTx623 and P721N (the progenitor of P721Q), and we have initiated crosses between P721Q and BTx623 to identify these modifiers.

## CONCLUSIONS and FUTURE DIRECTIONS

The inhibition of seed protein digestibility after wet cooking in sorghum is under genetic control. We have characterized multiple mutants that improve this digestibility and describe additional genetic factors that contribute to this improvement, that may provide important new targets for breeding efforts to improve food security. One of these, a novel, ankyrin repeat domain protein, HDG1, can be altered to provide better digestibility with more durable seed, and further investigation of this gene should reveal the mechanisms underlying these improvements and facilitate their optimization. Further investigation is also needed to understand the genetic modifiers in BTx623 that appear to enhance the protein digestibility of the kafirin signal peptide mutant SbEMS3324. The observation that at least one, and likely more, non-kafirin genes appear to be involved in this trait, and that mutations in them may have less adverse impacts on seed quality suggest protein digestibility is a valuable, complex trait to understand and improve. Our results provide two new avenues to explore and hold promise for breeding improved digestibility into elite, food grade lines that can contribute to food security in areas where sorghum is a staple crop.

## MATERIALS AND METHODS

### Plant material

Sorghum mutants SbEMS1227, SbEMS1613, and SbEMS3324 were identified in an EMS mutagenized population of BTx623 described previously(Krothapalli *et al.*, 2013; Addo-Quaye *et al.*, 2018). Seeds of the highly digestible, high lysine mutant, P721Q (Q for opaque endosperm) and its progenitor P721N (N for normal endosperm) were generously provided by Gebisa Ejeta.

### Development of mapping and validation populations

A mapping population of F_2:3_ families was generated for SbEMS3324 by first crossing it as a male onto the low digestibility homozygous *brown midrib 6* (*bmr6)* mutant SbEMS932. Similarly, a mapping population of F_2:3_ families was generated from SbEMS1613 by crossing it as a male onto SbEMS932. F1 seeds were generated in summer 2014 at the Purdue Agronomy Center for Research and Education (ACRE), West Lafayette (Indiana, USA). F1 seeds were planted during summer 2015, F1 plants identified as having white midribs, and these were self-fertilized to obtain F2 seeds. F2 plants were self-fertilized in summer 2016 at ACRE, and 455 SbEMS3324 and 506 SbEMS1613 F_2:3_ panicles obtained.

Validation populations were generated by crossing each of the highly digestible EMS mutants to low digestibility inbred lines and selecting F1 hybrid progeny. SbEMS1613 was crossed to CE151-262A, MR732, Tx430, and BTx623. SbEMS3324 was crossed to varieties Macia, TARG1, and MR732. F1 plants were selfed to produce panicles of F2 seed that were then planted panicle to row and selfed, with leaf tissue sampled from each F2 plant. The resulting F_2:3_ seed were assayed for digestibility and leaf tissue from plants with high digestibility (see below) were genotyped to search for shared mutations among progeny from crosses made with the mutant as a shared parent.

Thirty panicles from each generated population were tested for digestibility using triplicate pairs of seed as described below, and DNA extracted from leaves corresponding to individual, highly digestible progeny. PCR primers specific for the genes harboring candidate mutations were used to amplify either Sobic005G083340 or Sobic.005G189000 as appropriate, which were then Sanger sequenced using the PCR primers. Sequence alignment was then performed between the high and low digestibility F_2_ of the mapping populations and high digestibility F_3_ recombinants.

### Determination of protein digestibility

Protein digestibility was measured using a modified digestibility assay performed as follows:

On day one, 60 ± 2 mg of seeds were placed in a 2 mL reinforced screw-cap polypropylene tube with 6 ceramic beads (2.8 mm diameter) and 830 µL of double distilled water was added. Samples were ground in a BeadRuptor 24 (Omni International) at 6.5 m/sec for 60 s, then cooled at 4°C to prevent starch gelatinization. This grinding and cooling cycle was repeated three times or until the product was completely homogenized. Samples were then cooked into a microporridge for 20 min at 95°C in a VWR model 5420 rotating oven on maximum speed. The samples were left at room temperature to cool down and then vortexed one at a time to get a homogenous suspension. Using a wide-bore pipette tip, 300 µl of each sample were immediately put in two adjacent wells of a 96 deep well plate: one well to be digested with pepsin and the other one to be used as a control. The plate was then stored at 4 °C overnight.

The second day, samples were resuspended and 300 µL of a porcine pepsin solution (9 mg/mL of 3300 U/mg of pepsin (J.T Baker) in 0.2 M KH_2_PO_4,_ pH 2) was added to each sample to be digested, 300 µL of 0.2 M KH_2_PO_4_ pH 2 was added to each control well, the wells sealed with strip caps and the plate mixed by shaking. Digestion was performed in a rotating oven at 37 °C for 2 h. Digestion was stopped by adding 200 µL of 2 N NaOH to all 96 wells and the plates resealed. Samples were then centrifuged at 3700 rpm for 5 min. The supernatant was poured off and 500 µL of 0.1 M KH_2_PO_4_ pH 7 was added to each well to neutralize the pH, and plates vortexed for 5 minutes or until the pellet was resuspended. Samples were again centrifuged at 3700 rpm for 5 minutes, the supernatant poured off and each sample washed by adding 500 µL of double distilled water to each well, resealing the plate, and vortexing for 5 minutes or until the pellets were all resuspended. Samples were centrifuged at 3700 rpm for 5 minutes, the supernatant poured off and the pellets resuspended in 500 µL of Protein Extraction Buffer (1% SDS (w/v), 0.05% beta-mercaptoethanol (v/v), 0.0125 M sodium tetraborate solution, pH 10) and incubated at 25°C, 250 rpm for 1 h in racks with the tubes horizontal. Plates were centrifuged at 3700 rpm for 20 min. 20 µl of the supernatant were then transferred to a microtiter well in an optical flat-bottom plate containing 200 µL autoclaved distilled water. After all samples had been transferred, 50 µL of 72% trichloro acetic acid (w/v) was added to all samples to precipitate the remaining protein, the samples mixed manually, avoiding bubbles and the plate incubated at room temperature for 20 min before reading A_562_ using an EL × 800 UV spectrophotometer (BioTek Instruments, Inc.).

A560 values of each sample were used to calculate protein digestibility (PD) as follows:

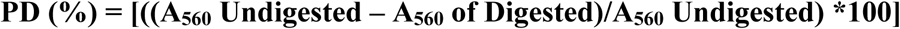

### Protein body morphology of sorghum mutants

Seeds were harvested 30 days after half bloom, fixed and sectioned by the Purdue Microscopy facility, following a previously described procedure (Oria *et al.*, 2000) and imaged using a Philips CM-100 transmission electron microscope (FEI Company). The microscope was operated at accelerating voltage of 100 kV with the following settings: spot 3,200 µm condenser aperture and 70 µm objective aperture. Images were taken using a Gatan digital camera.

### Determination of Protein, total lysine and tryptophan content

Five grams of seeds of P721Q, BTx623, SbEMS1613 and SbEMS3324 were analyzed for crude protein, total lysine and tryptophan content at the Agricultural Exeriment Station Chemical Laboratories at the University of Missouri. Tryptophan content was measured using the Enzymatic hydrolysis – colorimetric determination method (Draher and White, 2017). Total lysine was determined following the Association of Official Agricultural Chemists (AOAC) approved acid hydrolysis method. Crude protein content was obtained from the formula **Protein = %N * 6.25** where nitrogen was determined by the Kjeldhal method.

### Measuring seed diameter

Seed diameter of mutants and wild type parents was measured using a digital caliper. Five mature, dried seeds were randomly selected for each sample. Each measurement was taken transversely in the middle part of the seed, a mean value calculated and compared at p = 0.05 for all entries.

### Determination of seed hardness

Seed hardness measurements were conducted at the Purdue University Mechanical Engineering Department using an MTS 858 Mini Bionix instrument designed for static testing of material strength. Measurements were expressed as amount of force (in Newtons) needed to break a single seed fed to the instrument. Each genotype was measured twice, and an average hardness value was calculated.

### DNA extraction

Approximately 2 inches of the youngest fully expanded leaf was collected for each plant and placed in a marked envelope with one tablespoon of silica gel to allow leaves to dry; silica gel was recharged and changed frequently. Tissue samples were then bulked and DNA extracted using the CTAB method (Allen *et al.*, 2006): a bulk of 50 samples each from low digestibility plants and high digestibility plants for SbEMS1613; 30 plants for each bulk for SbEMS3324. The concentration and purity of the DNA was assayed on a Thermo Scientific Nanodrop 1000 Spectrophotometer and samples with A_260/280_ between 1.8 and 2.0 were considered pure. A total of 2 ng of DNA in a volume of 120 µL for each pool was used for whole genome sequencing at the Purdue Genomics Core Facility.

### DNA sequencing and detection of SNPs

Whole genome sequencing on pooled DNA was carried out using an Illumina Hiseq to generate 100 bp paired-end reads. Whole genome sequencing data of two of the lines (SbEMS1613 and SbEMS932) had been generated previously (Addo-Quaye *et al.*, 2018). Each sample or bulk received a unique barcode before being placed in the same flow cell lane.

After removal of the contigs, whole genome sequencing reads of each bulk were mapped against version 3.1.1 of sorghum reference genome (Paterson *et al.*, 2009) that was obtained online from Phytozome version 12.1.6 (Goodstein *et al.*, 2012). Alignment was achieved through the Burrows Wheeler Aligner (BWA) software package designed for alignment of short reads (Li and Durbin, 2009). SNPs were called using the command multiallelic-caller of SAMtools module version 1.4 (Li *et al.*, 2009). Only homozygous SNPs with a minimum quality of 20 and maximum depth of 250 were kept in Variant Call Format (VCF) files.

After obtaining the VCF files containing the filtered SNPs, all subsequent analyses were performed in R version 3.3 using the packages VariantAnnotation and Zoo. Allele frequencies were calculated and used to compute a rolling average difference between allele frequencies in the highly and lowly digestible groups for every 21 bp for SbEMS1613 and for every 11bp for SbEMS3324 mapping populations.

### SNP validation by Sanger sequencing and analysis of conserved regions

To verify results from bulked segregant analysis, DNA primers were identified using primer-BLAST (Ye *et al.*, 2012) based on the reference genome sequence region flanking the identified SNPs and were used to amplify targets in selected progeny of both mapping and validation populations. PCR amplification was performed as follows: 1 cycle of 98°C for 30 sec; 40 cycles of 10 sec denaturation at 98°C, 30 sec at the relevant primer hybridization temperature and extension at 72°C for 30 sec; and 1 cycle of final extension at 72°C for 5 min. Primer information is listed in Table 3. PCR products were gel purified and extracted from agarose using a Qiagen PCR purification kit. Quality of purified DNA was checked using a ThermoScientific NanoDrop spectrophtometer. DNA samples with A_260/A280_ between 1.8 and 2 were considered high quality and were Sanger sequenced. The resulting DNA sequences of parent, homozygous high digestibility and homozygous low digestibility progeny were aligned using BioEdit Sequence Alignment Editor version 7.2.6.

**Table 3.**
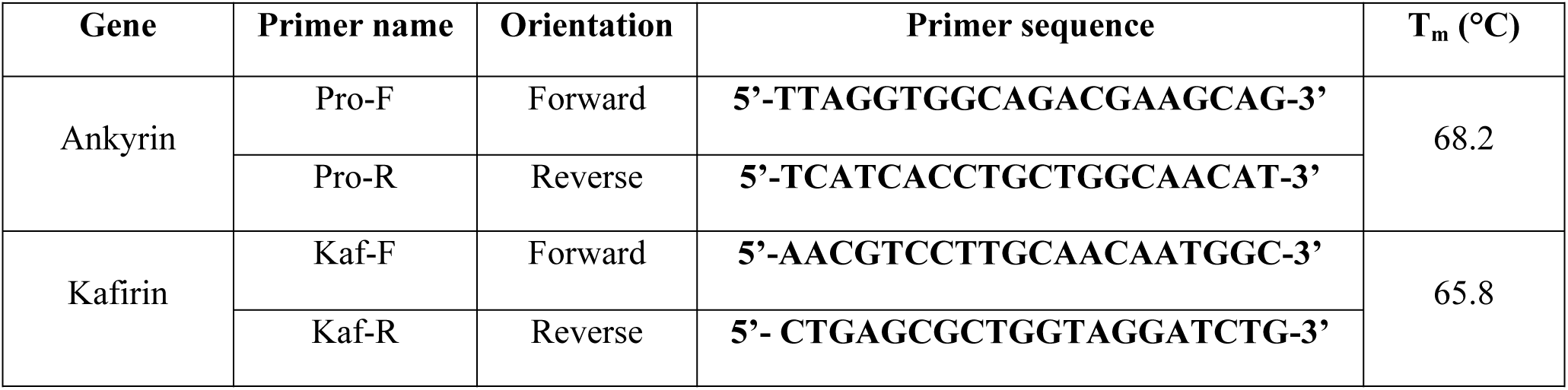
Primer information per gene

## Acknowledgments

The authors thank Jacqueline Anderson and Andrew Linvill for invaluable technical assistance. This research was made possible by the support of the Bill and Melinda Gates Foundation and support of the American people provided to the Feed the Future Innovation Lab for Collaborative Research on Sorghum and Millet through the United States Agency for International Development (USAID). The contents are the sole responsibility of the authors and do not necessarily reflect the views of USAID or the United States Government.

